# A comprehensive Arabidopsis transcription factor binding atlas reveals pervasive positional and syntactic organization of their DNA binding

**DOI:** 10.64898/2026.09.23.753179

**Authors:** Alice Jegou, Jérémy Lucas, Marianne Dreuillet, François Parcy, Romain Blanc-Mathieu

## Abstract

Transcription factor (TF) binding underlies gene regulation, yet the determinants of TF-DNA interactions across plant genomes remain incompletely understood. Here, we present a comprehensive atlas of TF binding in *Arabidopsis thaliana*, generated by reanalyzing 1,157 publicly available ChIP-seq and DAP/ampDAP-seq datasets using a unified processing framework. After stringent curation, 681 high-quality experiments were retained, providing binding information for 425 TFs across 42 families and 23 structural classes.

Using this resource, we show that TF binding predictability varies widely across TFs, with family identity explaining substantially more variation in predictive performance than structural class. Among the tested TF binding site (TFBS) prediction models, deep learning approaches achieved the highest overall predictive accuracy, while classical position weight matrices remained competitive and readily interpretable. Beyond motif recognition, we uncover widespread organizational principles of TF binding. TFBS positioning relative to transcription start sites is strongly family-dependent and more concentrated near promoters *in vivo*. Moreover, preferred spacing between homotypic TFBS is pervasive across TF families, indicating that binding-site syntax is a general organizational feature of TF binding in plants. Comparison of *in vivo* and *in vitro* binding profiles further identifies a shared core of sequence-driven binding alongside *in vivo*-enriched sites associated with non-canonical or composite sequence features.

Together, these results highlight the interplay between intrinsic DNA recognition, higher-order binding syntax, and cellular context in shaping TF binding landscapes, while providing a broadly useful resource for the plant community.

## Main

Transcription factors (TFs) are central regulators of gene expression, orchestrating developmental programs and responses to environmental cues by binding to specific DNA sequences and regulating nearby genes. Understanding how TFs recognize and bind their target sites across the genome remains a fundamental challenge in plant biology. While sequence-specific DNA recognition is often described by short motifs, accumulating evidence indicates that TF binding is shaped by a broader set of determinants, including nucleotide dependencies, DNA structural features, and the spatial organization of binding sites, such as preferred spacing, orientation, and positioning of homo- or heterotypic motifs. These intrinsic sequence features operate in conjunction with chromatin context and interactions with cofactors to define TF binding landscapes (Inukai et al. 2017; Lai et al. 2019; Rieu et al. 2023). Disentangling these contributions is essential for a comprehensive understanding of transcriptional regulation.

In addition, plant TFs comprise a highly diverse set of DNA-binding domain (DBD) architectures, recently classified into structural superclasses, classes, and evolutionarily related families according to their mode of DNA recognition in the Plant-TFClass resource (Blanc-Mathieu et al. 2024). Structural superclasses and classes describe increasingly refined modes of DNA recognition based on DBD architecture and DNA-contacting structural elements. Families, in contrast, represent evolutionarily related groups sharing common ancestry and generally similar DNA-binding preferences. While some DBD types are conserved across eukaryotes, others are specific to plants (e.g. LFY, TCP and GRAS domains) or have undergone substantial expansion in plant genomes (e.g. MADS-box and bHLH TFs) (De Mendoza et al. 2013; Wilhelmsson et al. 2017; Lai, Chahtane, et al. 2020). These structural and evolutionary distinctions are associated with diverse modes of DNA recognition, raising the question of how TF family and structural class contribute to the predictability and organization of TF binding across the genome.

Genome-wide TF-DNA binding assays, including chromatin immunoprecipitation followed by sequencing (ChIP-seq) and DNA affinity purification sequencing (DAP-seq), provide unprecedented genomic insights into TF binding landscapes in plants (Kaufmann et al. 2010; O’Malley et al. 2016; Song et al. 2016; Baumgart et al. 2025). ChIP-seq captures TF binding in its native chromatin context, reflecting the combined influence of sequence information, chromatin accessibility, and protein-protein interactions. In contrast, DAP-seq and ampDAP-seq, a variant in which DNA amplification removes methylation marks, measure intrinsic DNA-binding preferences in a controlled setting, largely independent of chromatin. These complementary approaches offer a powerful framework to disentangle sequence-driven and context-dependent components of TF binding (O’Malley et al. 2016; Lai et al. 2019). However, differences in experimental design, data processing, and analysis strategies have so far limited integrative analyses across datasets and prevented a unified view of TF binding landscapes (Ammari et al. 2026).

To predict TF binding, numerous computational models have been developed, ranging from classical position weight matrix (PWM)-based approaches to models incorporating nucleotide dependencies, DNA shape features, and deep learning architectures (Lai et al. 2019). While these models have demonstrated varying levels of success, their relative performance across TF families and experimental conditions remains incompletely characterized. Moreover, most studies focus on motif-level recognition, whereas higher-order organizational features of TF binding, such as the positioning of TF binding site (TFBS) relative to transcription start sites (TSS) or the spacing between homo- and heterotypic binding sites, have comparatively received little attention at the genome-wide scale in plants.

Indeed, studies of individual TF families have shown that binding-site syntax, including preferred inter-site distances and orientations, can reflect cooperative binding mechanisms, ranging from flexible TF-DNA binding geometries in Auxin Response Factors (ARFs) to higher-order assemblies on DNA in MADS-domain transcription factors (Boer et al. 2014; Lai, Vega-Léon, et al. 2021; Cancé et al. 2022). Whether such syntactic constraints represent a general property of plant TFs remains unclear. Likewise, the extent to which these features differ between *in vitro* and *in vivo* conditions, and how they can be integrated into sequence-based models of TF binding, is poorly understood.

Here, we address these questions by constructing TransAt, a comprehensive atlas of TF binding in *Arabidopsis thaliana*, through the uniform reanalysis of publicly available ChIP-seq and DAP/ampDAP-seq datasets. This resource integrates hundreds of experiments across diverse TF families, enabling a global and comparative view of TF-DNA interactions. Using this atlas, we benchmark multiple classes of TF binding site models and show that predictive performance is primarily determined by TF family rather than structural class, with deep learning approaches providing the highest overall predictive performance while classical PWMs remain competitive and readily interpretable.

Beyond motif recognition, we uncover widespread organizational principles of TF binding. TFBS positioning relative to TSS is strongly family-dependent and becomes more concentrated near promoter regions *in vivo*, consistent with the influence of cellular context. In addition, preferred spacing between homotypic TFBS is pervasive across TF families, indicating that binding site syntax, so far only described for a few TFs, is a widespread organizational feature of TF binding in plants. Finally, comparison of *in vivo* and *in vitro* binding profiles reveals a large shared core of sequence-driven binding, alongside a subset of in vivo-enriched genomic regions associated with non-canonical or composite sequence features.

Together, these results provide a unified framework for understanding TF binding in plants, highlighting the interplay between intrinsic DNA recognition, higher-order binding-site organization, and cellular context. The atlas and analyses presented here offer a resource for the plant community and a foundation for future studies aimed at deciphering the regulatory logic of gene expression.

### A comprehensive atlas of transcription factor binding sites in *Arabidopsis thaliana*

To enable a systematic analysis of TF binding in *Arabidopsis thaliana*, we assembled a comprehensive and uniformly processed collection of genome-wide TF-DNA binding datasets. We reprocessed 1,157 publicly available ChIP-seq (*in vivo*) and DAP/ampDAP-seq (*in vitro*) experiments using a unified analysis framework (<u>Methods</u>). After stringent quality filtering, requiring at least 600 high-confidence peaks (bound regions determined from derived-sequencing reads accumulation) and significant enrichment of a DNA motif known to be recognized by the assayed TF, 681 datasets (∼60%) were retained, comprising 110 *in vivo* and 571 *in vitro* experiments. This curated dataset provides high-confidence binding information for 425 distinct TFs. Because most TFs were assayed in only one experimental context, comparisons between *in vitro* and *in vivo* datasets are primarily performed at the TF-family level. Direct comparisons were possible for only 22 TFs represented by 75 datasets in both conditions.

To assess the extent to which the atlas captures TF diversity, we assembled and manually curated a set of 1,627 *Arabidopsis thaliana* TFs, which we classified into the 56 families defined in the Plant-TFClass resource (Blanc-Mathieu et al. 2024) (Supplementary Table 1). Although only 26% (425/1,627) of *Arabidopsis thaliana* TFs were represented by high-quality datasets after curation, these encompass 75% (42/56) of the TF families defined in Plant-TFClass, including 39 families represented by *in vitro* datasets and 24 by *in vivo* datasets. Representation remains uneven across families, ranging from ∼5% (1/21 TALE-type homeodomain TFs) to ∼71% (5/7 BES/BZRTFs) (Fig. 1a). Owing to their size, families such as ERF/DREB, MYB, NAC, and bZIP account for a substantial fraction of the datasets. Pairwise comparisons of genome-wide binding profiles revealed strong similarities among *in vitro* datasets for members of the same TF family, producing well-defined family-specific correlation clusters (Supplementary Fig. 1). In contrast, the *in vivo* correlation structure was markedly more heterogeneous (Supplementary Fig. 1). Although several well-represented regulators, including ELONGATED HYPOCOTYL 5 (HY5), E2FA, FRUITFULL (FUL), LEAFY (LFY), and SEPALLATA 3 (SEP3), formed coherent clusters, correlations were generally weaker and less clearly organized by TF family, consistent with the influence of chromatin accessibility, cofactor interactions and other regulatory constraints beyond intrinsic DNA-binding specificity.

**Fig. 1:**
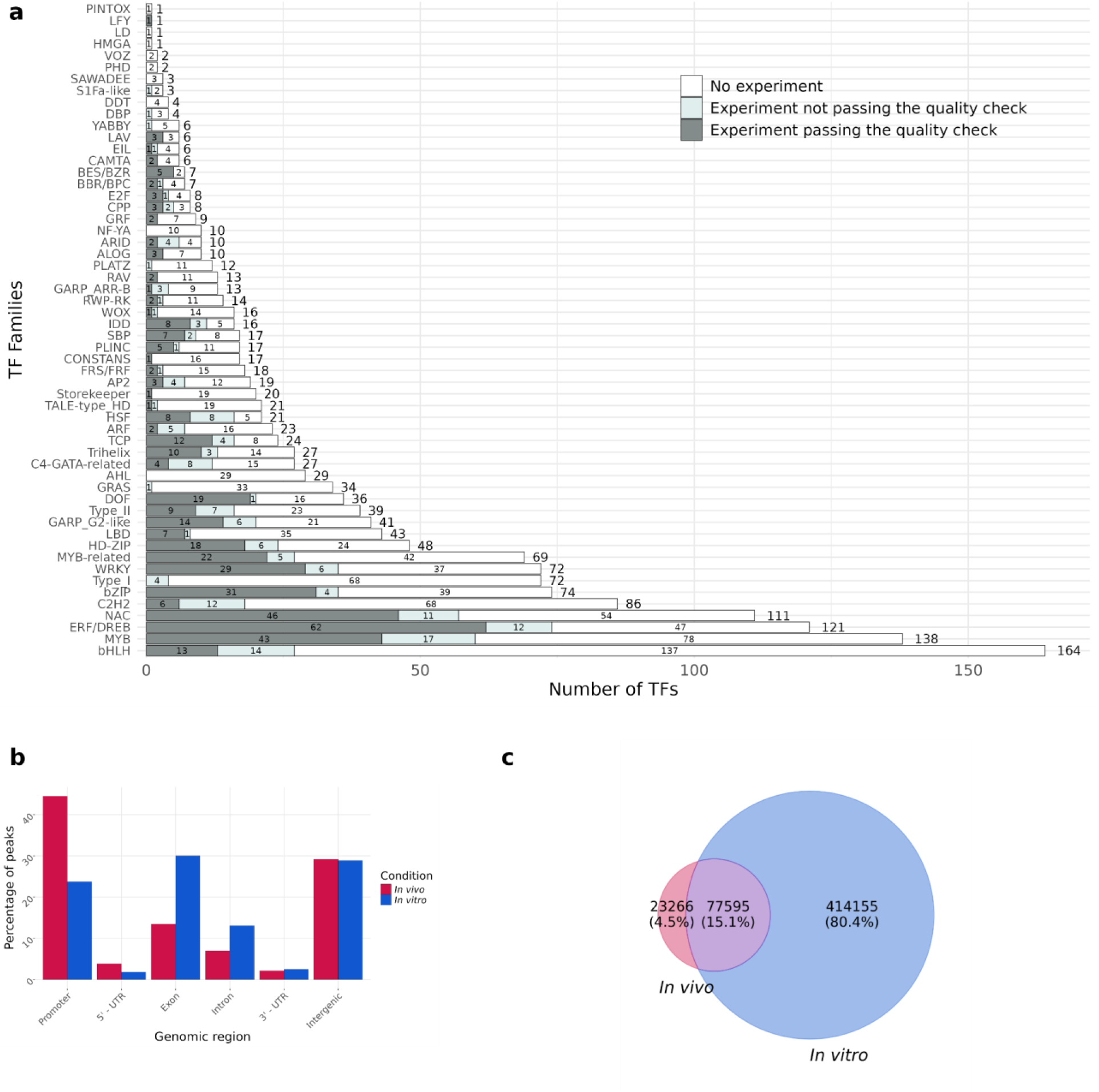
Composition and genomic distribution of the *Arabidopsis thaliana* transcription factor binding atlas. **a**, Distribution of genome-wide TF-DNA binding datasets across TF families. Bars indicate the number of TFs per family, colored by data availability: no experiments (white), experiments excluded after curation (light grey), and experiments retained for analysis (≥600 peaks and a validated DNA-binding motif; dark grey). **b**, Genomic distribution of peaks across annotated regions (promoters, UTRs, exons, introns, and intergenic regions) for in vivo (ChIP-seq) and in vitro (DAP/ampDAP-seq) datasets. **c**, Overlap of 515,016 merged, non-redundant peaks identified from in vivo (ChIP-seq) and in vitro (DAP/ampDAP-seq) datasets, showing shared and in vivo- or in vitro-specific binding events.

We next compared the genomic landscapes captured by *in vitro* and *in vivo* experiments. Across all datasets, *in vitro* peaks, resized to ±100 bp around their summit positions, covered 75% of the reference genome at depth ≥1, compared with 18% for *in vivo* datasets, reflecting both the broader binding landscapes typically observed in DAP/ampDAP-seq (Ammari et al. 2026) and the cumulative aggregation of hundreds of TF binding profiles rather than pervasive binding by individual TFs. ChIP-seq peaks were preferentially enriched in promoter regions (45% versus 24% in DAP/ampDAP-seq) and 5′ UTRs, but less frequently located within gene bodies than DAP/ampDAP-seq peaks, including both exons (13% versus 30%) and introns (7% versus 13%) (Fig. 1b). Intergenic regions were similarly represented in both assay types (∼29%). These differences likely reflect the influence of chromatin context and other regulatory constraints on TF binding *in vivo*, whereas DAP/ampDAP-seq primarily captures intrinsic DNA-binding preferences.

To provide a unified reference framework for downstream analyses, we merged peaks across all experiments to define a non-redundant set of TF-bound regions. This procedure identified 515,016 regions, the majority originating from *in vitro* datasets (Fig. 1c). The most closely related resources are the Plant Cistrome (O’Malley et al. 2016), which contains exclusively DAP/ampDAP-seq datasets, and ReMap2022 (Hammal et al. 2022), which integrates binding profiles from diverse classes of transcriptional and chromatin regulators and defines non-redundant peak sets independently for each factor. In contrast, our atlas focuses specifically on TFs and provides experiment-specific peak sets, quantitative binding profiles, and a unified catalogue of non-redundant TF-bound regions across all datasets.

### Transcription factor binding model performance is driven by family-specific features

To assess how accurately TF binding can be predicted from DNA sequence, we benchmarked five classes of TFBS models using the atlas described above. Model performance was evaluated for both *in vitro* and *in vivo* datasets using the area under the receiver operating characteristic curve (AUROC) on balanced test sets (<u>Methods</u>). The benchmark includes five TFBS modeling approaches: position weight matrices (PWMs) (Stormo et al. 1982), transcription factor flexible models (TFFMs) (Mathelier and Wasserman 2013), DNA shape-augmented models (Mathelier et al. 2016), k-mer set memory (KSM) models (Guo et al. 2018), and SeqConv convolutional neural network (CNN) models (Shen et al. 2021). PWM, TFFM, DNA shape-augmented and KSM models all infer TF binding from motif-centric representations, whereas SeqConv learns predictive sequence features directly from raw DNA sequences. Given its widespread use, PWM served as the reference model throughout the analyses.

Overall, all models achieved good predictive performance, with AUROC values generally exceeding 0.7 and consistently higher performance *in vitro* than *in vivo* (Fig. 2). We first examined how performance varied across TF families before comparing the relative performance of the different modeling approaches.

**Fig. 2:**
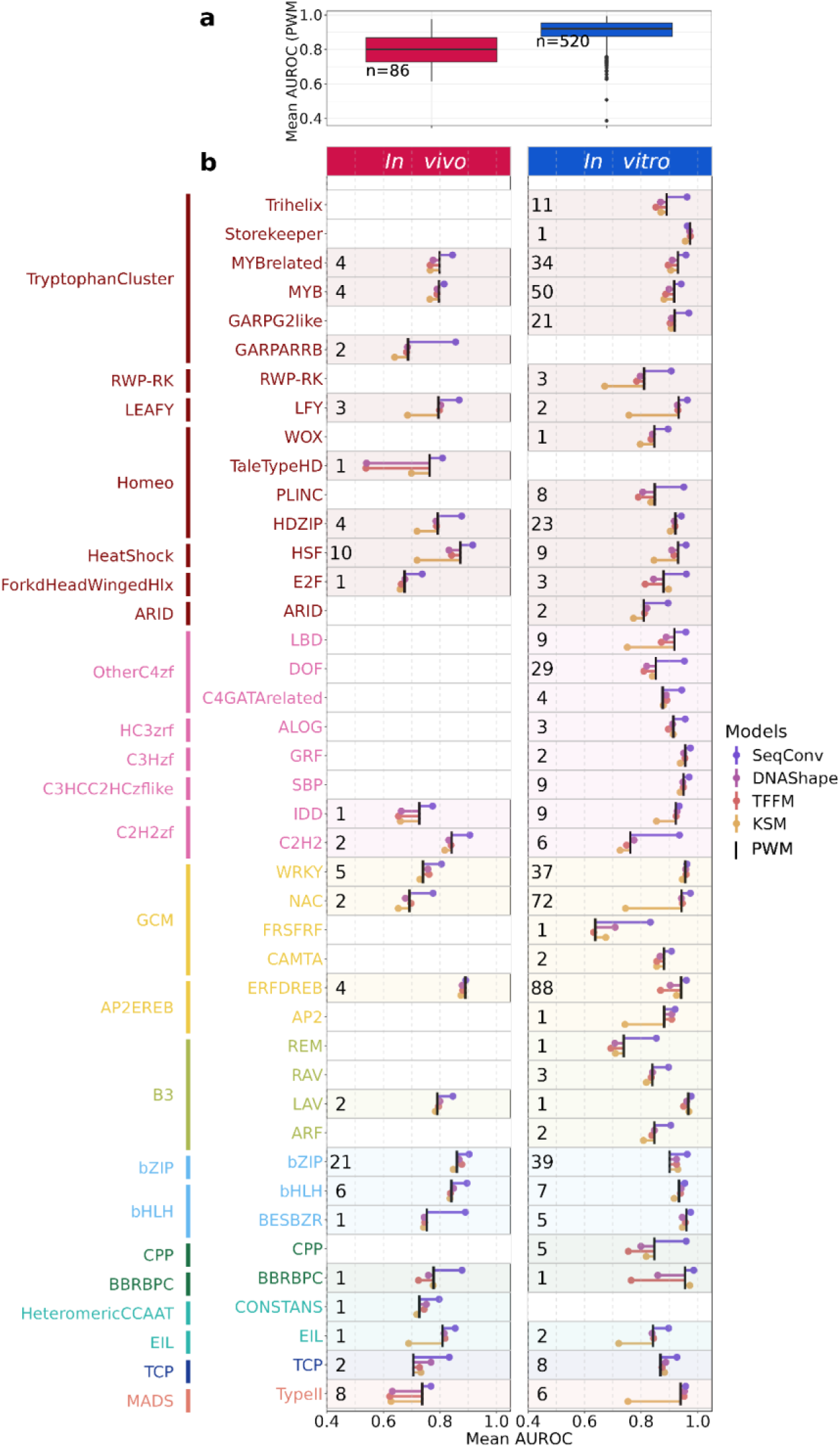
Performance of transcription factor binding models across families *in vitro* and *in vivo*. **a,** Distribution of PWM performance across all TF binding datasets, shown as AUROC values for *in vivo* (ChIP-seq) and *in vitro* (DAP/ampDAP-seq) datasets. **b,** Mean AUROC per TF family for *in vivo* (left) and *in vitro* (right) datasets. Families are colored by structural superclass and ordered by class along the y-axis; numbers indicate the number of samples per family. PWM is shown as the reference model (black tick), and alternative models (TFFM, KSM, DNAshape, SeqConv) are represented as lollipops indicating their mean AUROC relative to PWM. Values higher than the PWM reference indicate improved predictive performance, whereas lower values indicate reduced performance. Values were logit-transformed prior to family-level averaging and back-transformed for visualization.

We found that the model predictive performance varied markedly between TF families. Several families, including BES/BZR and WRKY, achieved near-perfect discrimination (AUROC ∼ 1), whereas a small subset of datasets exhibited PWM AUROC values below 0.7 (Fig. 2a). To investigate the origin of this variability, we examined the relationship between PWM performance and motif information content. AUROC increased with motif information content, indicating that TFs recognizing highly specific DNA motifs are generally easier to predict because their binding sites are more readily distinguished from genomic background sequences (Supplementary Fig. 2). Additionally, variance partitioning using a mixed-effects model revealed that TF family explains a substantial fraction of PWMs performance variance (∼34%), whereas structural class contributes only marginally (∼3%) (Supplementary Fig. 3). Similar variance partitioning patterns were observed for TFFM and DNA shape-augmented models, consistent with their shared motif-centric framework.

We next compared the relative performance of the different TFBS models. SeqConv consistently outperformed or matched the other approaches across nearly all TF families, although improvements over PWM were generally modest for TFs whose motifs were already highly informative. In contrast to motif-based approaches, SeqConv showed a marked reduction in family-level performance variance (∼9%), accompanied by a narrower distribution of AUROC values across TF families (Supplementary Fig. 3). This suggests that CNN-based models capture sequence features beyond canonical motifs, resulting in more homogeneous predictive performance across TF families.

To determine whether the relative performance of the different models changed when the evaluation was restricted to the highest-confidence predictions, we also assessed model performance in the low false-positive-rate region of the ROC curve (FPR ≤ 1%), which is particularly relevant for genome-wide TFBS prediction (Supplementary Fig. 4). Although SeqConv achieved the highest overall AUROC, its relative performance decreased *in vivo* when the evaluation was restricted to the low-FPR region. One possible explanation is that CNNs exploit sequence features associated with TF occupancy beyond the core binding motif. While these additional features improve overall discrimination, they may reduce the relative contribution of strong canonical motifs, leading to lower performance among the highest-confidence predictions. The fact that this effect was observed only *in vivo* further suggests that these additional sequence features may reflect biological determinants of TF occupancy that are absent from *in vitro* binding assays.

Together, these results indicate that TF binding predictability is strongly influenced by family-specific sequence properties rather than broad structural TF classes. This variability has important practical implications, as the reliability of TFBS predictions can differ substantially among TFs and should therefore be considered when using these models to identify candidate binding sites. Deep learning models achieved the highest overall predictive performance, with substantial improvements over conventional motif-based approaches for many TF families, although the magnitude of this gain varied across families and experimental contexts. The remaining variability in predictive performance further indicates that important determinants of TF binding are incompletely captured by current sequence-based models, particularly under cellular conditions.

### Family-dependent positioning of transcription factor binding sites around transcription start sites

Previous studies have reported that *Arabidopsis thaliana* TFs often exhibit preferential binding either upstream or downstream of transcription start sites (TSSs), frequently in a family-dependent manner (Yu et al. 2016; Voichek et al. 2024). Orientation biases of non-palindromic TFBS have also been reported (Lis and Walther 2016). However, systematic comparisons of TFBSs positioning and orientation across TF families, and between *in vivo* and *in vitro* conditions, are limited. To address this gap, we quantified the enrichment of PWM-predicted TFBSs within TF-bound regions overlapping a ±1 kb window around TSSs (<u>Methods</u>) and summarized family-level positional and orientation preferences (Fig. 3).

**Fig. 3:**
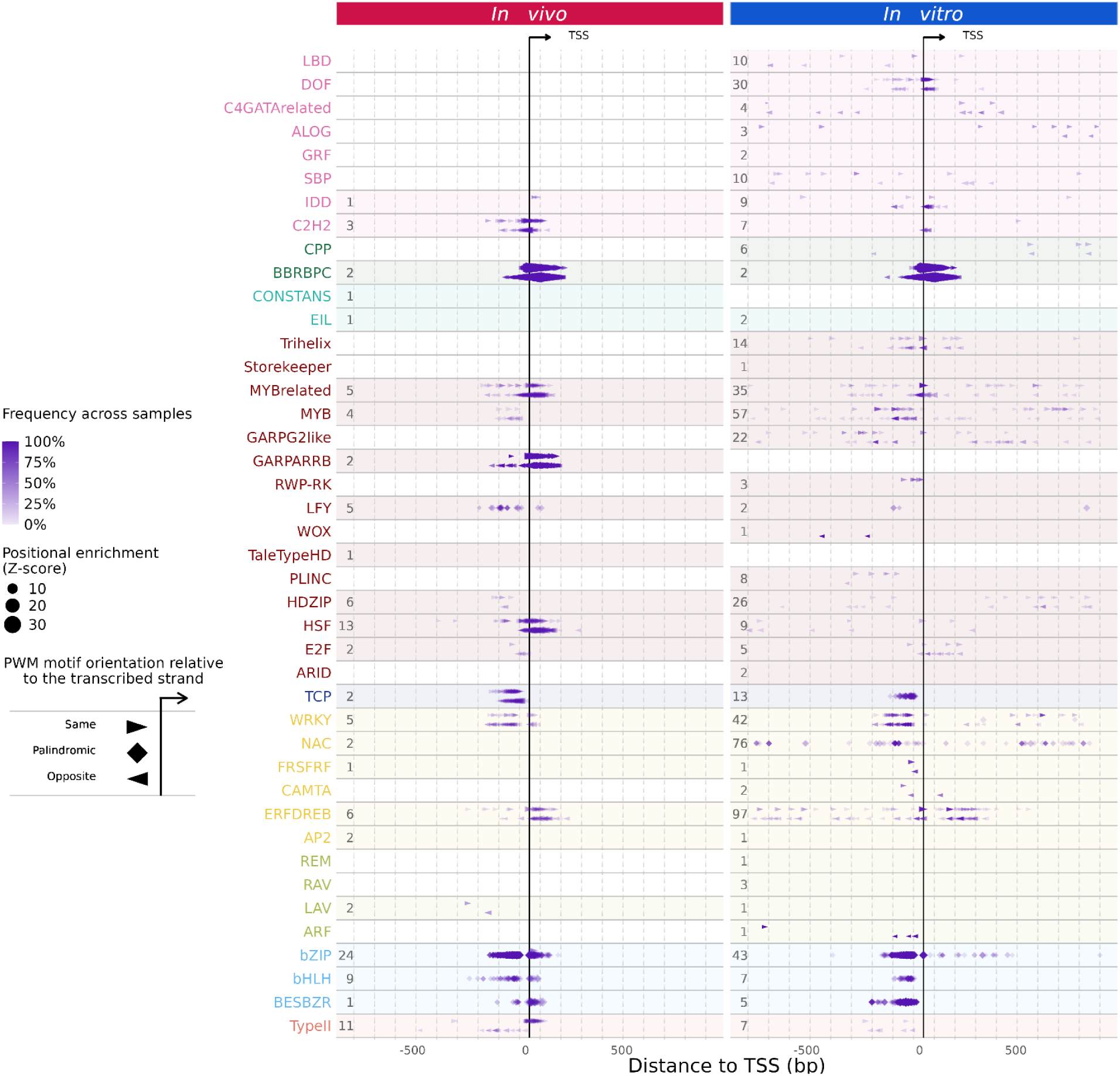
Positional preferences of transcription factor binding sites relative to transcription start sites. Enriched positions of transcription factor binding sites (TFBS) identified within experimental peaks relative to transcription start sites (TSS) across TF families. Left, *in vivo* (ChIP-seq); right, *in vitro* (DAP/ampDAP-seq). The x-axis indicates the distance to the TSS (-1 kb to +1 kb), and the y-axis lists TF families grouped by structural superclass, with the number of samples indicated. Symbols mark significantly enriched TFBS positions (median Z-score). Triangles represent non-palindromic TFBS and indicate motif orientation relative to the TSS, shown on separate sub-rows. Diamonds represent palindromic TFBS. Symbol size reflects enrichment strength (average median Z-score across TFs within a family), and color intensity indicates the proportion of samples exhibiting the positional preference.

TFBSs exhibited clear family-specific positional preferences relative to TSSs. Significantly enriched TFBS-TSS distances frequently displayed a positional bias relative to the TSS, with a significant upstream or downstream bias detected in 13 of 31 families *in vitro* (42%) and 14 of 18 families *in vivo* (78%) (Supplementary Table 2). Orientation biases were less frequent, occurring in 3 of 25 families *in vitro* (12%) and 7 of 15 families *in vivo* (47%) with at least one enriched distance associated with an orientable motif (Supplementary Table 3). Several positional preferences were conserved across conditions, including downstream enrichment for BBRBPC and ERF/DREB and upstream enrichment for bZIP, TCP and WRKY. Rather than extending uniformly across promoters, these orientation biases are typically confined to discrete regions relative to the TSS. For example, Type II MADS and GARP/ARR-B TFBSs exhibit localized enrichments over narrow distance ranges.

Experimental context reveals distinct positional binding landscapes. *In vitro*, enriched TFBSs are distributed over a broad range around TSSs, often extending across the full ±1 kb window. In contrast, TFBSs detected in ChIP-seq peaks are markedly concentrated near TSSs, typically within ±200 bp, consistent across most TF families. This shift toward promoter-proximal enrichment likely reflects the combined influence of chromatin accessibility and other cellular constraints acting *in vivo*. Notably, for several TF families (e.g., HSF, WRKY, ERF/DREB), *in vivo* conditions are associated with a marked reinforcement of promoter-proximal enrichment near TSSs. In some cases, weak or diffuse positional preferences observed *in vitro* become sharply concentrated near the TSSs *in vivo,* whereas more distal enrichments are reduced or absent, suggesting that cellular context preferentially constrains TF binding toward promoter-proximal regions.

Together, these results indicate that TFBS positioning relative to TSSs is a widespread, family-dependent feature. Although orientation biases are less common than positional biases, they are observed in a few TF families. The enrichment of TFBSs near TSSs *in vivo* suggests that promoter architecture and cellular context contribute to shaping TF binding beyond intrinsic DNA sequence preferences.

### Widespread homotypic TFBS syntax across transcription factor families

In plants, preferred binding-site organization between homotypic TFBSs, including defined spacing and orientation constraints, has been documented for specific TF families such as ARFs and MADS-domain proteins, where these configurations support cooperative binding and higher-order complex assembly on DNA (Stigliani et al. 2019; Lai, Stigliani, et al. 2020). More broadly, clusters of homotypic TFBSs have been observed in plant regulatory regions and have been proposed to contribute to TFs occupancy and regulatory activity (Barah et al. 2016). However, the extent to which such organizational features represent a general property of TF binding across plant TF families remains largely unexplored.

To systematically identify potential syntax rules across TF families, we searched TFBSs within peak sequences using PWM-based scanning and quantified their pairwise distances and orientations (<u>Methods</u>), enabling a family-level analysis of homotypic spacing and orientation preferences. For TFs known to bind DNA as obligate dimers (e.g., LFY, bZIP, NAC family members), dimeric PWMs were used. Indeed, unlike TFs such as ARF5, which can bind pairs of monomeric auxin response cis-elements (AuxREs) separated by alternative spacings (Boer et al. 2014), such TFs recognize composite motifs with a fixed internal organization. Consequently, the detected configurations reflect arrangements between dimeric binding units and provide candidate configurations for higher-order assemblies.

Unexpectedly, enriched homotypic TFBS configurations were detected across a broad range of TF families (Fig. 4), indicating that preferred spacing and orientation between TFBSs are not restricted to a few well-studied families but represent a common feature of plant TF binding. Two recurring patterns emerge across TF families. First, several families display clusters of neighboring preferred spacings, comprising multiple closely related configurations within a narrow distance range. Examples include E2F, FRS/FRF and C4-GATA-related TFs. Second, some families exhibit periodic spacing arrangements, characterized by recurring configurations separated by regular intervals, as observed for LAV, WOX, LBD and ERF/DREB families. Both patterns indicate that TF binding can be accommodated with multiple preferred configurations rather than a single fixed arrangement.

**Fig. 4:**
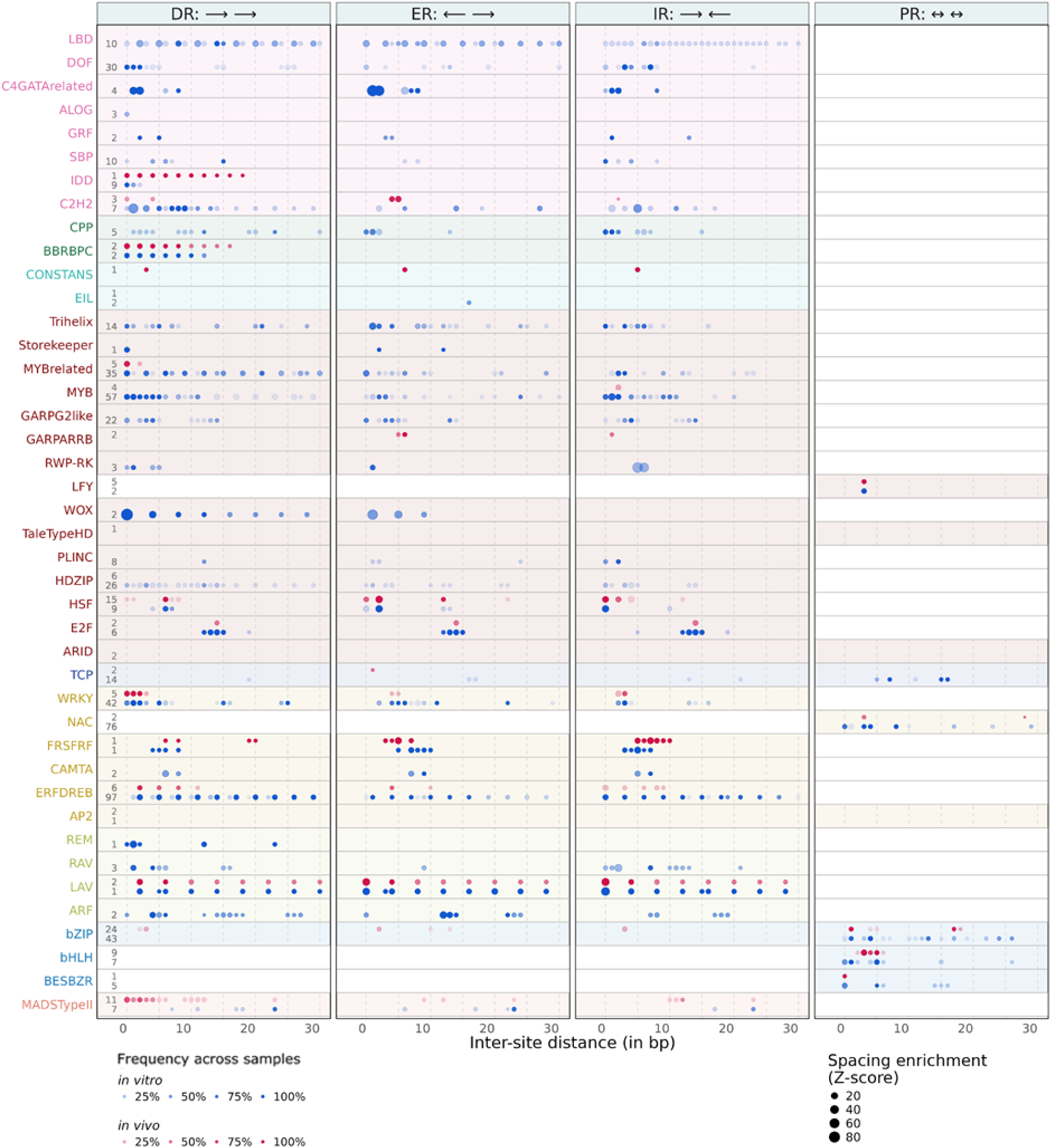
Enriched distances and orientations between pairs of transcription factor binding sites (TFBS) across TF families. Panels show direct (DR), everted (ER), inverted (IR), and palindromic (PR) repeat configurations. The x-axis indicates inter-site distance (0-30 bp), and the y-axis lists TF families grouped by structural superclass. Dots mark significantly enriched TFBS arrangements detected within peak regions: blue for *in vitro* (DAP/ampDAP-seq) and red for *in vivo* (ChIP-seq). Dot size reflects enrichment strength (average median Z-score across TFs within a family), and color intensity indicates the proportion of samples exhibiting the corresponding configuration. Numbers next to each TF family denote the number of *in vitro* and *in vivo* samples. Enrichment was assessed for distances up to 70 bp; only distances ≤30 bp are shown for clarity, as few significant spacings were detected beyond this range.

The conservation of these configurations also varies among families. In some cases, the same spacing patterns are recovered across most TFs within a family. For example, all E2F family members exhibit clusters of preferred spacings in the 12-15 bp range *in vitro*, suggesting a shared binding syntax. In other families, dominant configurations are accompanied by lower-frequency variants restricted to a subset of TFs, indicating family-wide trends combined with TF-specific preferences.

Comparison between *in vitro* and *in vivo* datasets revealed substantial conservation of spacing preferences across the two conditions. At the family level, similar spacing patterns were recovered in both datasets for several TF families, including LAV, ERF/DREB and BBRBPC, suggesting that many syntax rules are largely independent of the experimental context (Fig. 4). Although these family-level comparisons are based on only partially overlapping sets of TFs, direct comparison of the 22 TFs represented by at least one *in vivo* and one *in vitro* dataset supported this observation. Fourteen TFs displayed congruent spacing patterns, with most enrichments detected *in vivo* also being recovered *in vitro* (Supplementary Table 4). Among the remaining TFs, the lack of agreement generally resulted from the absence of significant spacing enrichments in one condition rather than from conflicting spacing preferences. Non-overlapping enrichment patterns were observed only for SEP3 and HY5. Together, these results suggest that a substantial fraction of homotypic syntax reflects intrinsic properties of TF-DNA recognition that are maintained across experimental conditions.

Despite this overall conservation, condition-specific enrichments were also observed. For instance, type II MADS TFs SEP3, APETALA1 (AP1) and FUL display a series of closely spaced configurations *in vivo* that are not detected *in vitro*. These short-range enrichments likely reflect local clustering of CArG boxes, the canonical binding sites of MADS-box transcription factors, rather than the long-range organization associated with floral quartet complexes formed by SEP3 and AGAMOUS (AG) (Lai, Stigliani, et al. 2020). Although not reaching our significance criteria, AG ChIP-seq datasets from floral tissues also showed enrichments near the ∼37 bp and ∼47 bp spacings previously associated with MADS-domain tetrameric complexes (Lai, Stigliani, et al. 2020). Conversely, some spacing patterns detected *in vitro* (e.g., in MYB, MYB-related, and C2H2 families) are not recovered *in vivo*. This may reflect the broader binding landscape accessible in the absence of chromatin constraints, although the limited representation of these TFs in *in vivo* datasets may also contribute.

Together, these results indicate that homotypic TFBS syntax is a pervasive and family-dependent feature of TF binding in *Arabidopsis thaliana*. The widespread occurrence of preferred spacing and orientation constraints across TF families suggests that information relevant to TF recognition is frequently encoded not only in motif sequence, but also in the relative arrangement of binding sites, highlighting binding-site syntax as an additional layer of binding and possibly regulatory information. Several non-exclusive molecular mechanisms may underlie the diversity of binding-site organizations observed across TF families. First, spacing flexibility may arise from geometrical tolerance within the TF-DNA complex, either through conformational flexibility of the protein assembly itself or through local deformation of the bound DNA, as illustrated by ARF5, which can bind inverted AuxRE repeats separated by either 7 or 8 bp (Boer et al., 2014). Conversely, some TF-DNA complexes can impose stringent spacing requirements: WUSCHEL binds cooperatively to tandem TGAA repeats through DNA-mediated homodimerization, whereas altering the orientation of the repeats or introducing a spacer strongly reduces binding affinity (Sloan et al. 2020). Second, alternative oligomerization interfaces could generate distinct binding geometries. Although such mechanisms have not yet been reported for plant TFs, the bacterial regulator ArdK forms a symmetric dimer on inverted repeats and an asymmetric dimer on direct repeats, thereby recognizing different motif organizations through alternative protein-protein interfaces (Fernandez-Lopez et al. 2022). Finally, cooperativity may be mediated in part by the DNA itself, whereby local sequence context and DNA shape facilitate binding across a broader range of spacings (Jolma et al. 2013; Sloan et al. 2020). The relative contribution of protein-protein interactions and DNA-mediated cooperativity is likely context-dependent and remains difficult to disentangle. Importantly, several studies suggest that these syntax rules can have functional consequences. Specific AuxRE configurations are enriched in promoters of auxin-responsive genes (Stigliani et al. 2019), and synthetic promoter assays have demonstrated that distinct AuxRE architectures generate different transcriptional outputs and expression patterns *in planta* (Martin-Arevalillo et al. 2025). Similarly, changes in protein identity among MADS-box transcription factors are associated with shifts in preferred CArG-box spacing and corresponding changes in developmental function (Lai, Vega-Léon, et al. 2021). While our analysis identifies numerous candidate syntax rules across *Arabidopsis* TF families, further work will be required to determine which configurations contribute directly to cooperative complex formation and transcriptional regulation.

### Cellular context modulates transcription factor binding beyond intrinsic DNA recognition

ChIP-seq and DAP/ampDAP-seq provide complementary views of TF binding. Whereas DAP/ampDAP-seq primarily captures intrinsic DNA-binding specificity, ChIP-seq additionally reflects the influence of chromatin organization, cofactors and other cellular determinants of TF occupancy. To investigate how these factors shape TF binding profiles, we compared 75 matched ChIP-seq and DAP/ampDAP-seq experiments representing 22 TFs.

Across all comparisons, distributions of log2 fold changes in normalized peak coverage were centered near zero (Fig. 5a), indicating that a large fraction of binding regions exhibit similar occupancy *in vitro* and *in vivo*. This shared binding landscape is consistent with binding events primarily determined by intrinsic sequence features, including canonical TFBSs and their local organization, rather than by chromatin or other cellular factors. These results further support the ability of DAP/ampDAP-seq to capture the intrinsic DNA-binding preferences of TFs.

**Fig. 5:**
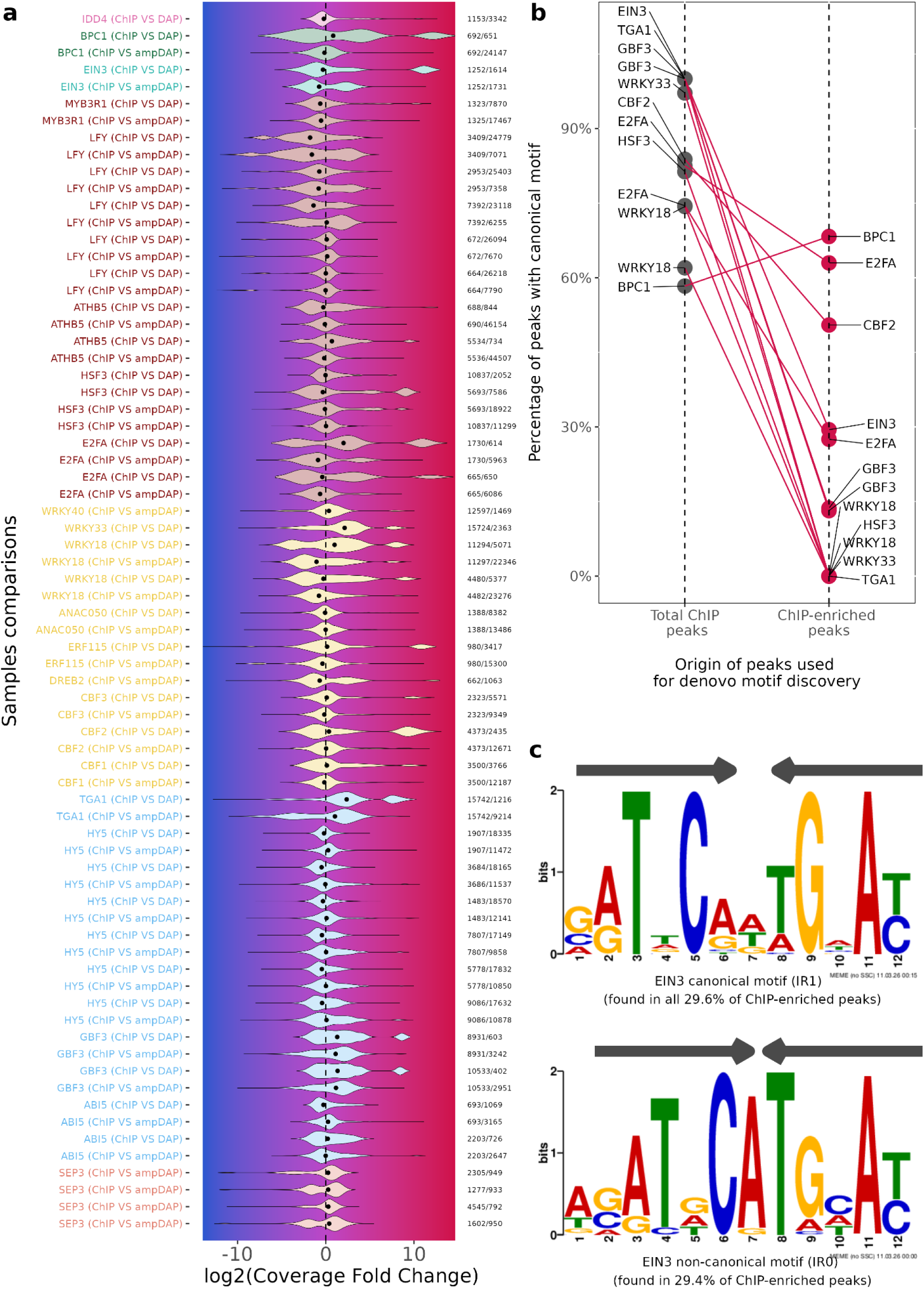
Cellular context modulates transcription factor binding beyond intrinsic DNA recognition. **a**, Comparison of normalized peak coverage between matched ChIP-seq and DAP/ampDAP-seq datasets for TFs represented in both experimental systems. Violin plots show the distribution of log2 fold changes in normalized peak coverage (ChIP/DAP) calculated over the combined set of shared and condition-specific genomic regions used for pairwise comparison. Positive values indicate higher occupancy *in vivo*, whereas negative values indicate higher occupancy *in vitro*. The vertical dashed line denotes equal occupancy between conditions. Numbers indicate the number of peaks *in vitro* or *in vivo*. Comparisons are ordered and coloured according to TF structural superclass. **b**, Proportion of peaks containing the canonical TF family motif among the highest occupied ChIP-seq peaks (grey) and among manually defined *in vivo*-enriched peak subsets (red). Motifs were identified by *de novo* motif discovery using MEME. **c**, Example of a non-canonical motif identified in *in vivo*-enriched EIN3 binding regions. The canonical motif derived from the highest occupied ChIP-seq peaks is shown above, and the *in vivo*-enriched motif below. Binding of EIN3 to the canonical motif was confirmed by electrophoretic mobility shift assay (Supplementary Fig. 5), supporting its direct recognition *in vitro*. Grey arrows indicate the two inverted half-sites, which are separated by one nucleotide in the canonical motif (IR1) but are directly adjacent (IR0) in the *in vivo*-enriched motif.

Despite this substantial shared component, most TFs displayed asymmetric coverage distributions, revealing subsets of regions preferentially occupied in one experimental context. For several TFs, distributions extend towards negative log2 fold change values, corresponding to regions with lower occupancy *in vivo* than *in vitro*. Such regions likely represent binding sites where chromatin accessibility restricts TF binding, as previously reported for LFY, whose *in vivo*-depleted binding sites are enriched in closed chromatin states (Lai, Blanc-Mathieu, et al. 2021). Conversely, several TFs exhibited distinct populations of peaks with markedly higher occupancy *in vivo* than *in vitro*, forming secondary density maxima towards positive log2 fold change values. These *in vivo*-enriched regions were observed for members of the BBR/BPC, EIL, HSF, E2F, WRKY, ERF/DREB and bZIP families, and likely reflect TF binding promoted by chromatin accessibility, heteromeric protein complexes or other regulatory mechanisms that are absent from *in vitro* assays.

To characterize these *in vivo*-enriched regions, we performed exploratory motif analyses. Across multiple TFs, they contained a lower proportion of canonical TFBSs than the highest occupied ChIP-seq peaks (Fig. 5b). De novo motif discovery instead identified a diverse collection of sequence features, including motifs recognized by other TF families, non-canonical motif configurations and low-complexity sequence elements, consistent with previous observations that non-target TF motifs are common in ChIP-seq datasets (Worsley Hunt and Wasserman 2014).

For example, *in vivo*-enriched EIN3 peaks contained a non-canonical motif composed of two adjacent inverted repeats (IR0), whereas the strongest non-enriched ChIP-seq peaks only contained the canonical configuration in which the two inverted repeats are separated by a single nucleotide (IR1) (Fig. 5c). Although no single alternative motif architecture emerged across TFs, these observations suggest that context-dependent TF occupancy frequently involves combinatorial sequence features extending beyond canonical DNA-binding motifs.

Together, these analyses indicate that TF binding landscapes comprise a large sequence-driven component shared between *in vitro* and *in vivo* conditions, underscoring the ability of DAP/ampDAP-seq to capture intrinsic DNA-binding specificity. Superimposed on this core binding landscape is a smaller but likely biologically important set of context-dependent binding events shaped by chromatin state, cofactors and higher-order TF complexes. These regions provide a valuable framework for investigating the molecular mechanisms that modulate TF occupancy beyond sequence-encoded DNA recognition.

## Conclusion

Although TF-DNA interactions have been extensively studied, many aspects of the TF binding specificity and organization remain poorly understood, particularly in plants. In this study, we present TransAt, a comprehensive atlas of TF binding in *Arabidopsis thaliana*, generated through the systematic reanalysis of publicly available ChIP-seq and DAP/ampDAP-seq datasets using a unified processing framework. We leverage this resource to investigate the determinants of TF binding specificity and binding-site organization. This resource provides high-confidence binding information for hundreds of TFs across diverse families and experimental conditions, offering a global view of TF-DNA interactions in plants.

Our analyses reveal that TF binding specificity is primarily shaped by family-level sequence features rather than by the structural class of DNA-binding domains. While deep learning models achieve the highest overall predictive performance, classical PWM-based approaches remain robust and interpretable, capturing much of the intrinsic sequence specificity. However, predictive performance varies widely among TFs, and much of this variation cannot be explained by TF family or structural class alone. This indicates that important determinants of TF binding remain incompletely captured by current models, potentially including motif syntax, alternative binding configurations, or other sequence features beyond canonical motifs. These findings highlight both the strengths and limitations of current TFBS models in describing TF-DNA interactions, particularly in cellular contexts.

Beyond motif recognition, we uncover widespread organizational principles of TF binding sites. TFBS positioning relative to TSS is strongly family-dependent and differs markedly between *in vitro* and *in vivo* conditions, with promoter-proximal enrichment observed *in vivo*. This pattern is consistent with a recent work suggesting that TSS-proximal regions enriched in TFBSs may preferentially support accessible and dynamic regulatory activity, whereas distal elements may contribute to more stable regulatory programs (Morales-Cruz et al. 2026). In addition, we demonstrate that preferred spacing between homotypic TFBSs is a pervasive feature across TF families, extending well beyond previously characterized cases. These spacing patterns likely reflect constraints imposed by cooperative binding and higher-order complex formation on DNA. Importantly, the candidate spacing configurations identified here can help refine the annotation of functional regulatory sites. For example, distinct AuxRE architectures have been shown to generate different transcriptional outputs *in planta*, illustrating how TFBS syntax can encode regulatory information beyond the core binding motif (Martin-Arevalillo et al. 2025). These configurations may also generate testable hypotheses regarding TFs oligomer architecture, which could guide future structural and biochemical studies.

Finally, comparison of *in vivo* and *in vitro* binding profiles revealed that TF occupancy comprises a large sequence-driven core captured by DAP/ampDAP-seq together with a smaller set of context-dependent binding events observed only in cellular environments. These regions are associated with non-canonical or composite sequence features, suggesting that chromatin state, cofactors and cooperative DNA recognition expand the regulatory landscape beyond intrinsic DNA-binding specificity. Such mechanisms are exemplified by the LFY-UNUSUAL FLORAL ORGANS complex, which recognizes composite DNA motifs comprising a weaker canonical LFY binding site together with additional sequence features, thereby expanding the repertoire of bound genomic regions (Rieu et al. 2023).

Together, our results support a model in which TF binding is governed by a combination of intrinsic sequence recognition and higher-order regulatory syntax, modulated by cellular context. The TransAt resource and the analyses presented here provide a framework for investigating these mechanisms and for further dissecting the rules governing transcriptional regulation in plants. TransAt could also provide a valuable training resource for emerging deep-learning approaches aimed at deciphering TF binding specificity and regulatory syntax from genomic sequence (Peleke et al. 2026).

## Methods

### Transcription factors classification

All analyses were organized according to the Plant-TFClass framework, which groups plant TFs into superclasses based on DNA-binding domain (DBD) structure, further subdivided into structural classes and evolutionarily related families. Proteins (TFs and transcriptional regulators) were collected from multiple databases, manually curated to retain the ones that belong to families with evidence for specific DNA binding, and assigned to one of the 56 TF families defined in Plant-TFClass. Structural superclasses were used for color coding in figures.

### Data acquisition

Genome-wide TF-DNA binding datasets for *Arabidopsis thaliana* were collected from public repositories. Most DAP-seq and ampDAP-seq datasets were obtained from (O’Malley et al. 2016). Additional ChIP-seq and DAP-seq datasets were retrieved from ChIP-Hub (Fu et al. 2022), the Gene Expression Omnibus (GEO), and through manual curation of the literature. Queries to GEO were performed using combinations of TF names and relevant assay keywords. Selected experiments and specific parameters used for their processing are reported in Supplementary Table 5.

### Read mapping and peak calling

Raw sequencing datasets were reprocessed using a standardized pipeline adapted from our former studies (Lai, Stigliani, et al. 2020; Rieu et al. 2023). Sequence reads were downloaded from the NCBI Sequence Read Archive using the SRA Toolkit (v3.0.0), quality assessed with FastQC (v0.11.7), and trimmed of adaptors using NGmerge (v0.2_dev) (Gaspar 2018). Reads were aligned to the *Arabidopsis thaliana* TAIR10 reference genome (Lamesch et al. 2012) using Bowtie2 (v2.5.4) (Langmead and Salzberg 2012). Only primary alignments were retained, and reads with more than two mismatches or mapping quality ≤30 were discarded. PCR duplicates were removed using the MACS3 (v3.0.3) (Zhang et al. 2008) filterdup function, retaining at most one fragment per genomic position (--keep-dup 1).

Peak calling was performed independently for each biological replicate using MACS3 (v3.0.3) with the corresponding control library whenever available. Peaks were identified using a q-value threshold of 0.05, local background estimation (slocal = 500 bp; llocal = 5 kb), and summit detection enabled. Additional filtering steps removed peaks overlapping a custom blacklist derived from genomic regions showing reproducible enrichment across independent control datasets, as well as peaks showing less than threefold enrichment over the corresponding control signal (factork = 3). Reproducible peaks across all replicates were identified using MSPC (v5.4.0) (Jalili et al. 2016) with weak and strong p-value thresholds of 1 × 10⁻⁴ and 1 × 10⁻⁸, respectively, and only MSPC-supported consensus peaks were retained for downstream analyses. Consensus peak summits were refined by comparing summit positions among replicates, and final peak coordinates were centered on the consensus summit and resized to a fixed width of 200 bp. These parameters defined the gold-standard peak set.

To maximize dataset retention while maintaining reproducibility, two additional peak-calling stringency levels were defined. A silver standard (llocal = 2.5 kb, factork = 2.5, q-value = 0.2, MSPC weak = 1 × 10⁻⁴, MSPC strong = 1 × 10⁻⁸) and a bronze standard (llocal = 2.5 kb, factork = 2, q-value = 0.5, MSPC weak = 1 × 10⁻³, MSPC strong = 1 × 10⁻⁷) were used when the gold standard yielded fewer than 600 MSPC-supported consensus peaks, a threshold chosen to ensure sufficient data for reliable TFBS model inference and subsequent analyses. Silver and bronze standards were applied only to experiments for which at least two biological replicates were available.

### Motif discovery and dataset curation

For each dataset, peaks were ranked according to normalized read coverage maxima and the 600 highest peaks were used for *de novo* motif discovery with MEME (v4.12.0) (Bailey et al. 2015). Five motifs were requested for each dataset using parameters specified in Supplementary Table 5. Datasets were subsequently subjected to manual curation. Motifs were considered informative when they corresponded to known family-specific DNA-binding preferences, as assessed using JASPAR reference motifs (Rauluseviciute et al. 2024), or, in the absence of prior information, when they exhibited strong central enrichment within peaks, as measured by CentriMo (Bailey et al. 2015), and were detected in more than one-third of the training sequences. Only datasets containing at least 600 consensus peaks under one of the peak-calling standards (gold, silver, or bronze) and an informative motif were retained for further analyses.

### TFBS model construction

The manually curated MEME motif was converted into a position frequency matrix (PFM), which served as the basis for all motif-based models. Position weight matrices (PWMs) were generated directly from the curated PFMs. Transcription factor flexible models (TFFMs) and DNA shape-augmented models (Mathelier and Wasserman 2013; Mathelier et al. 2016) were subsequently trained using the corresponding PWM as input. In addition to PWM-based approaches, k-mer set memory (KSM) models (Guo et al. 2018) and convolutional neural network models (SeqConv; (Shen et al. 2021) were generated for each dataset. KSMs were computed using KMAC (v3.4) with kmer size between 5 and 25 bp. SeqConv was selected as a representative sequence-based deep-learning model because it was previously validated on large-scale *Arabidopsis* TF binding datasets. Model-specific parameters are provided in Supplementary Table 5.

### TFBS model benchmarking

Model performance was evaluated only for datasets containing at least 1,000 consensus peaks. For motif-based models (PWM, TFFM, DNA shape, and KSM), the 600 highest-confidence peaks were used for model construction and the remaining peaks were reserved for evaluation, thereby ensuring complete separation between training and testing sequences. Consequently, each dataset contained at least 400 positive test sequences. For SeqConv, 80% of peaks were randomly assigned to the training set and the remaining 20% to the test set, corresponding to a minimum of 800 training and 200 test sequences per dataset. This protocol differs from that used for the motif-based models but was adopted to accommodate the larger training requirements of deep learning approaches.

Model performance was assessed using receiver operating characteristic (ROC) curves on balanced test sets comprising positive sequences (held-out peaks) and an equal number of matched negative sequences. Negative sequences were generated as previously described (Rieu et al. 2023) by sampling genomic regions not bound by the corresponding TF while matching positive sequences for genomic annotation and GC content. For PWM, TFFM, DNA shape, and KSM models, each sequence was assigned the score of its highest-scoring motif occurrence. For SeqConv, scores corresponded to the predicted binding probabilities returned by the neural network. Model performance was quantified using the overall area under the ROC curve (AUROC) and the partial AUROC (pAUROC) over the false-positive rate interval [0,0.1], thereby emphasizing performance at low false-positive rates. To facilitate comparisons across models, pAUROC values were standardized following (McClish 1989) such that a random classifier scores 0.5 and a perfect classifier scores 1.

### TFBS models variance partitioning analysis

To assess the relative contribution of TF family and structural class to model performance, variance partitioning was performed using a linear mixed-effects model on AUROC values obtained from *in vitro* datasets. The TF family was modeled as a random effect nested within a structural class. Variance components corresponding to TF family, structural class, and residual variance were estimated from the fitted model, implemented in Python using the statsmodels mixed-effects framework with restricted maximum likelihood estimation. Analyses were restricted to TF families represented by at least three TFs and to structural classes containing at least two families. When multiple *in vitro* datasets were available for a given TF, only the sample with the highest AUC was retained to avoid over-representation.

### Homotypic TFBS spacing for syntax analysis

For each dataset, peak sequences were scanned using the corresponding PWM. Consequently, identified TFBS represent model-derived predictions rather than direct measurements of TF occupancy. Subsequences scoring above a specified PWM threshold were considered candidate TFBS. For each pair of TFBS within a peak, the inter-site distance (up to a maximum of 70-bp) and relative orientation (direct repeat, inverted repeat, or everted repeat) were computed as previously described (Stigliani et al. 2019). Inter-site distance was defined as the number of base pairs separating the edges of the two TFBS (edges are defined as left and right offsets relative to the PWM positions in Supplementary Table 5).

For each orientation class, the frequency of every spacing configuration was compared with the empirical distribution of all observed spacings within the dataset. Enrichment was quantified using a robust Z-score statistic based on the median and median absolute deviation (MAD) of the empirical spacing distribution.

Spacing configurations were considered significantly enriched when they satisfied all of the following criteria: (i) Bonferroni-adjusted p-value < 0.05, (ii) fold enrichment greater than two relative to the median background frequency, and (iii) reproducibility across at least two of three PWM score thresholds. PWM thresholds corresponded to the 99.9th, 99.7th, and 99.5th percentiles of PWM scores obtained from genome-wide scans (Supplementary Table 5).

Significant spacing configurations were defined as preferred spacings and represent enriched combinations of distance and orientation between homotypic TFBS.

### TFBS positioning and orientation relative to transcription start sites

To analyze TFBS positioning relative to transcription start sites (TSSs), peaks overlapping a ±1,000 bp window around annotated TAIR10 TSSs were selected. As in the homotypic spacing analysis, TFBS were identified by scanning peak sequences with the corresponding PWM and therefore represent model-derived predictions rather than direct measurements of TF occupancy.

Enrichment of TFBS positions relative to the TSS was quantified using the same robust median-based Z-score framework employed for homotypic spacing analysis. Position-specific enrichments were identified by comparing the observed frequency of TFBS at each position to the empirical distribution of TFBS positions within the dataset.

To assess positional and orientation biases, significantly enriched TFBS-TSS positions were classified according to their location upstream or downstream of the TSS and according to whether the corresponding TFBS was oriented in the same or opposite direction as the transcribed strand. For each TF family, the number of significantly enriched positions in each category was compared to the expected 1:1 ratio using two-sided exact binomial tests. Resulting p-values were corrected for multiple testing using the Benjamini-Hochberg procedure, and adjusted p-values < 0.05 were considered significant.

### Comparison of *in vivo* and *in vitro* binding profiles

For TFs represented by both ChIP-seq and DAP/ampDAP-seq datasets, peaks were resized to 200 bp centered on their summits. Peaks from paired experiments were considered to represent the same binding event when their overlap exceeded 80%. Shared binding regions were defined as the intersection of overlapping peaks, whereas non-overlapping peaks were retained as condition-specific regions. Read coverage was quantified over the resulting set of shared and condition-specific regions and normalized by the total number of reads mapped within peaks. When multiple biological replicates were available, normalized coverage values were averaged across replicates. Normalized peak coverage was then compared between ChIP-seq and DAP/ampDAP-seq experiments by calculating the log2 fold change in coverage.

### Electrophoretic Mobility Shift Assay (EMSA)

Candidate motifs identified during manual curation were evaluated by EMSA using EIN3, TCP4 and AtSTKL2. Coding sequences were synthesized (ShineGene Inc.), cloned into a 5×Myc-tag pTNT expression vector by Gibson Assembly using gene-specific primers (Supplementary Table 6), verified by Sanger sequencing, and expressed in vitro using the TNT® SP6 High Yield Wheat Germ Protein Expression System (Promega). Briefly, 2 μg of plasmid DNA were added to 24 μL of TNT® extract in a final reaction volume of 40 μL and incubated for 2 h at 25 °C. Protein expression was verified by western blot using the iBlot™ 2 Dry Blotting System (Thermo Fisher Scientific). Three double-stranded DNA probes were designed for each TF (Supplementary Table 7), fluorescently labelled with Cy5 using the Klenow fragment (New England Biolabs), and incubated with 5 μL of TNT® protein extract for 30 min on ice in binding buffer (40 mM HEPES-KOH pH 7.9, 100 mM KCl, 200 mM Tris-HCl pH 8.0 and 2.5% glycerol). DNA–protein complexes were resolved on 2% native agarose gels in 0.5× Tris–borate–EDTA (TBE) buffer at 4 °C and visualized using an ImageQuant 800 imaging system (GE Healthcare), as described previously (Martin-Arevalillo et al. 2025).

### Use of generative artificial intelligence

Le Chat (Mistral AI) and ChatGPT (OpenAI) were used during manuscript preparation to assist with language editing, sentence reformulation, and improving the clarity and readability of the text, as well as with the writing, modification, and debugging of analysis code. All AI-assisted code and scientific content were reviewed and validated by the authors, who take full responsibility for the analyses, interpretations, conclusions, and final manuscript.

## Data and code availability

All processed peak (BED) and coverage (BigWig) files, as well as TFBS models generated in this study and the Snakemake (7.32.4) (Mölder et al. 2021) workflow used to process data are publicly available as TransAt (Transcription Factor Binding Atlas for *Arabidopsis thaliana*) via Zenodo (https://doi.org/10.5281/zenodo.20556976). Raw sequencing data were obtained from public repositories (GEO/SRA), and reprocessed using a standardized pipeline available on GitHub ( https://github.com/Bioinfo-LPCV-RDF/TF_genomic_analysis_v2.git). The webpage of the structural classification Plant-TFClass is available at https://bioinfo-lpcv-rdf.github.io/Plant-TF-page/. The correspondence was made at the TF family level with the TAPscan database (Petroll et al. 2025) (https://tapscan.plantcode.cup.uni-freiburg.de/). The list of the 1,627 *Arabidopsis thaliana* transcription factors assigned to the plant-TFClass families is available in Supplementary Table 1. Complete lists of TFBS-TSS positioning preferences and homotypic TFBS spacing configurations identified for each dataset, together with their associated enrichment scores, are provided as Supplementary Tables 8 and Supplementary Table 9, respectively.

## Acknowledgments

We thank Renaud Dumas, Gabriel Krouk, Anthony Mathelier and Emmanuel Thévenon for their advice and helpful discussions, and Loïc David, Dipika Patel and Laura Turchi for their contributions to data collection and bioinformatics during the initial stages of the project. We are grateful to the Rensing laboratory for linking Plant-TFClass into the TAPscan database. We acknowledge the computing infrastructure at CEA Grenoble for providing the computational resources used in this study.

This work was supported by the GRAL LabEx (ANR-10-LABX-49-01), financed within the University Grenoble Alpes Graduate School (Écoles Universitaires de Recherche) CBH-EUR-GS (ANR-17-EURE-0003), and by the French National Research Agency through the “Investissements d’avenir” program (ANR-15-IDEX-02) to R.B.M., the ANR JCJC project TRANSINET (ANR-23-CE20-0027) to R.B.M., and the ANR project BEFLORE (ANR-21-CE20-0024) to F.P.

## Authors contribution

R.B.M and F.P conceived and supervised the project. A.J, J.L and R.B.M performed data collection, processing and downstream analyses. M.D performed the EMSA experiments. R.B.M, A.J and F.P wrote the manuscript with contributions from all authors.

## Supplementary information

**Supplementary Fig. 1:**
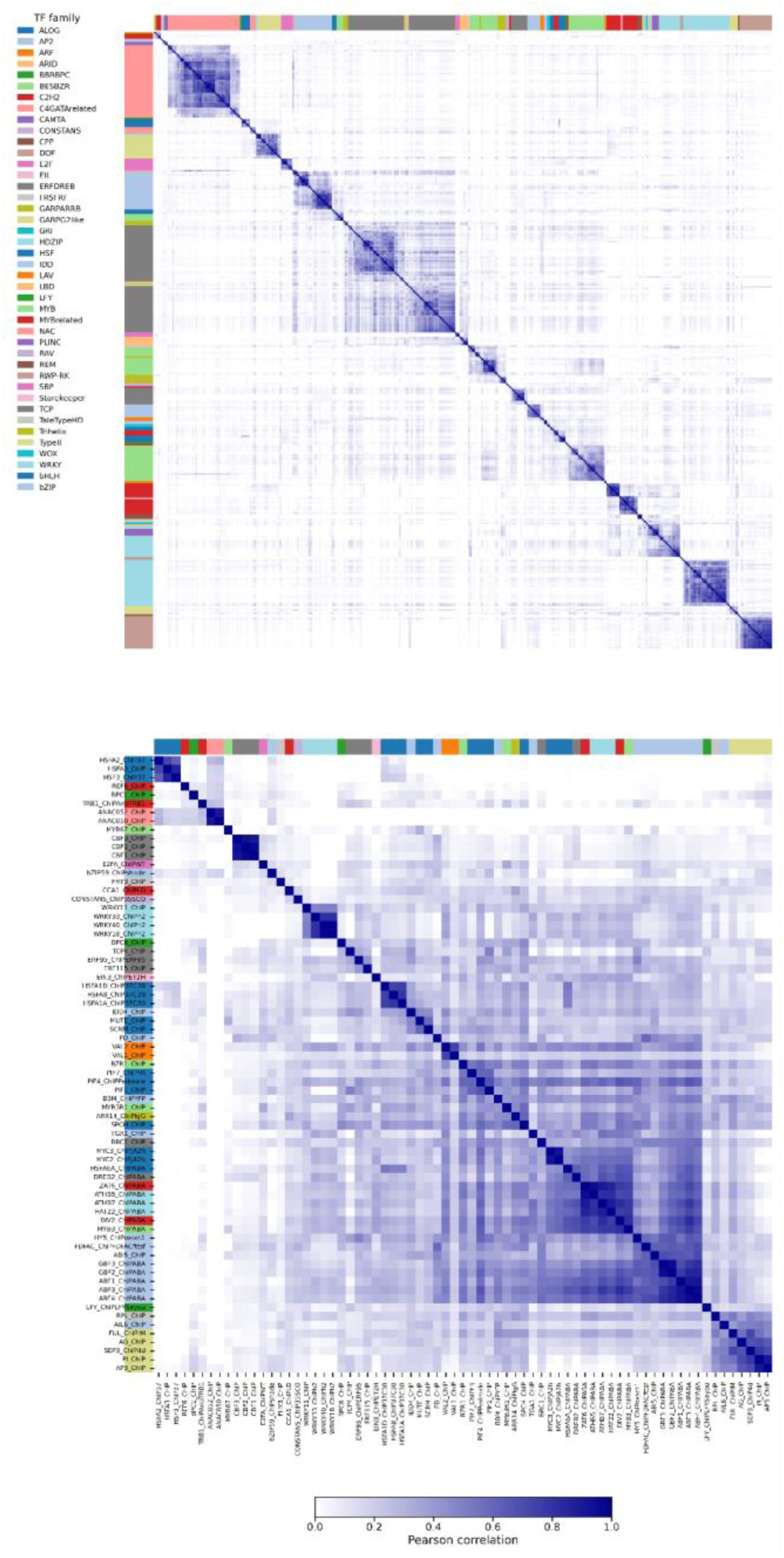
Pairwise correlation of transcription factor binding profiles across experiments. Heatmaps show pairwise Pearson correlation coefficients between genome-wide TF binding profiles from *in vivo* (left, ChIP-seq) and *in vitro* (right, DAP/ampDAP-seq) experiments. For each dataset, normalized read coverage was quantified over the corresponding set of merged peak regions, and Pearson correlation coefficients were calculated between all pairs of experiments. TF families are indicated by the colored bars.

**Supplementary Fig. 2:**
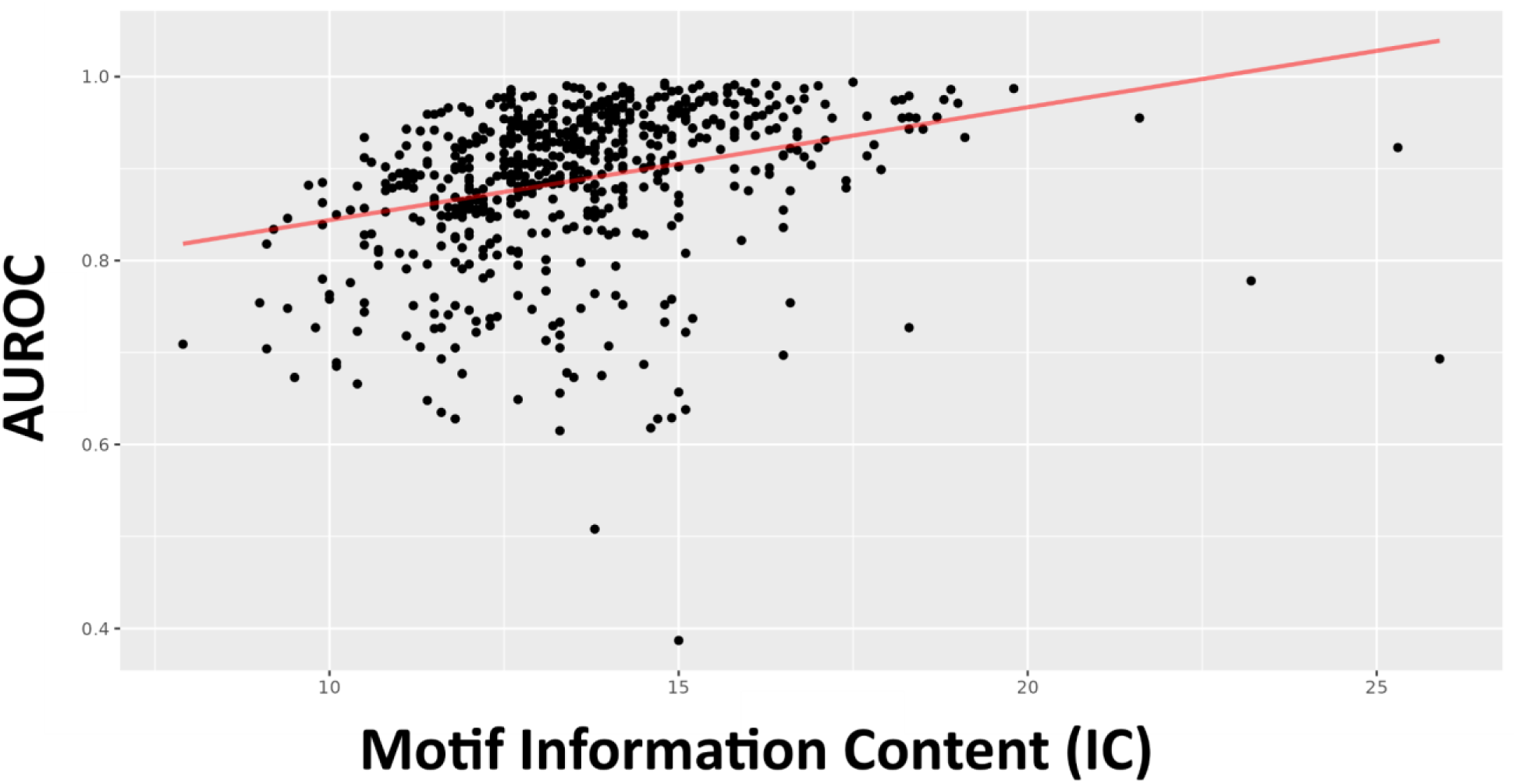
Relationship between motif information content and PWM predictive performance. PWM predictive performance (AUROC) plotted against motif information content (IC) for all TF binding datasets included in the study. Each point represents one experiment. The red line indicates the least-squares linear regression fit (R² = 0.09, *P* = 4.8e-14).

**Supplementary Fig. 3:**
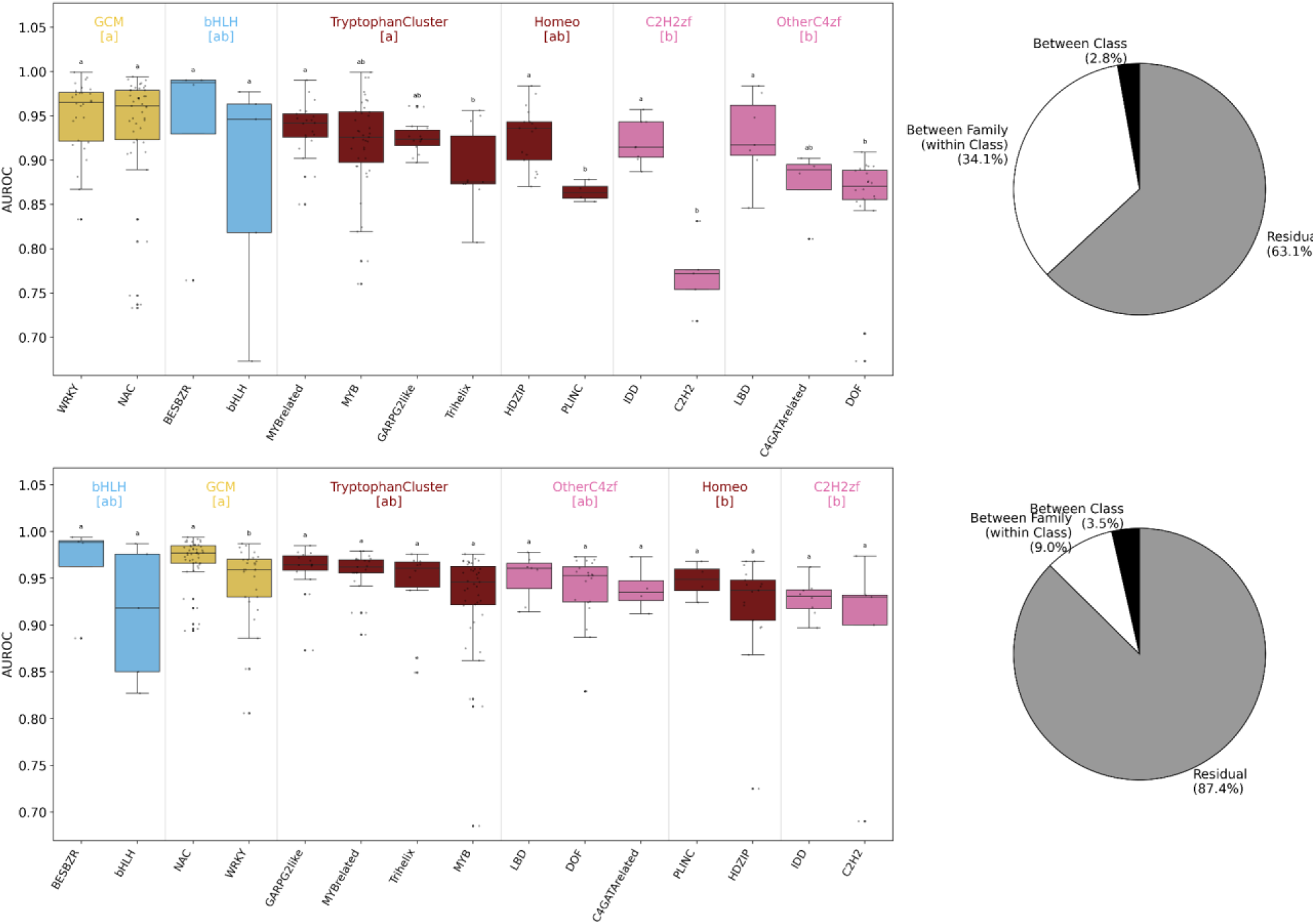
Variance partitioning of TF binding model performance across transcription factor families and structural classes. Nested distributions of AUROC values (left) are shown for PWM (top) and SeqConv (bottom) models across transcription factor (TF) families grouped within structural classes (*in vitro* datasets only, *n* = 231 TFs, distributed into 15 families and 6 classes). Boxplots represent AUC distributions for individual TF families nested within structural classes. Statistical differences between families were assessed using Tukey’s post hoc tests. Variance partitioning (right) was performed using a mixed-effects model to estimate the contribution of TF family (nested within structural class), structural class, and residual variance to model performance. Results are shown as variance components for PWM and SeqConv models.

**Supplementary Fig. 4:**
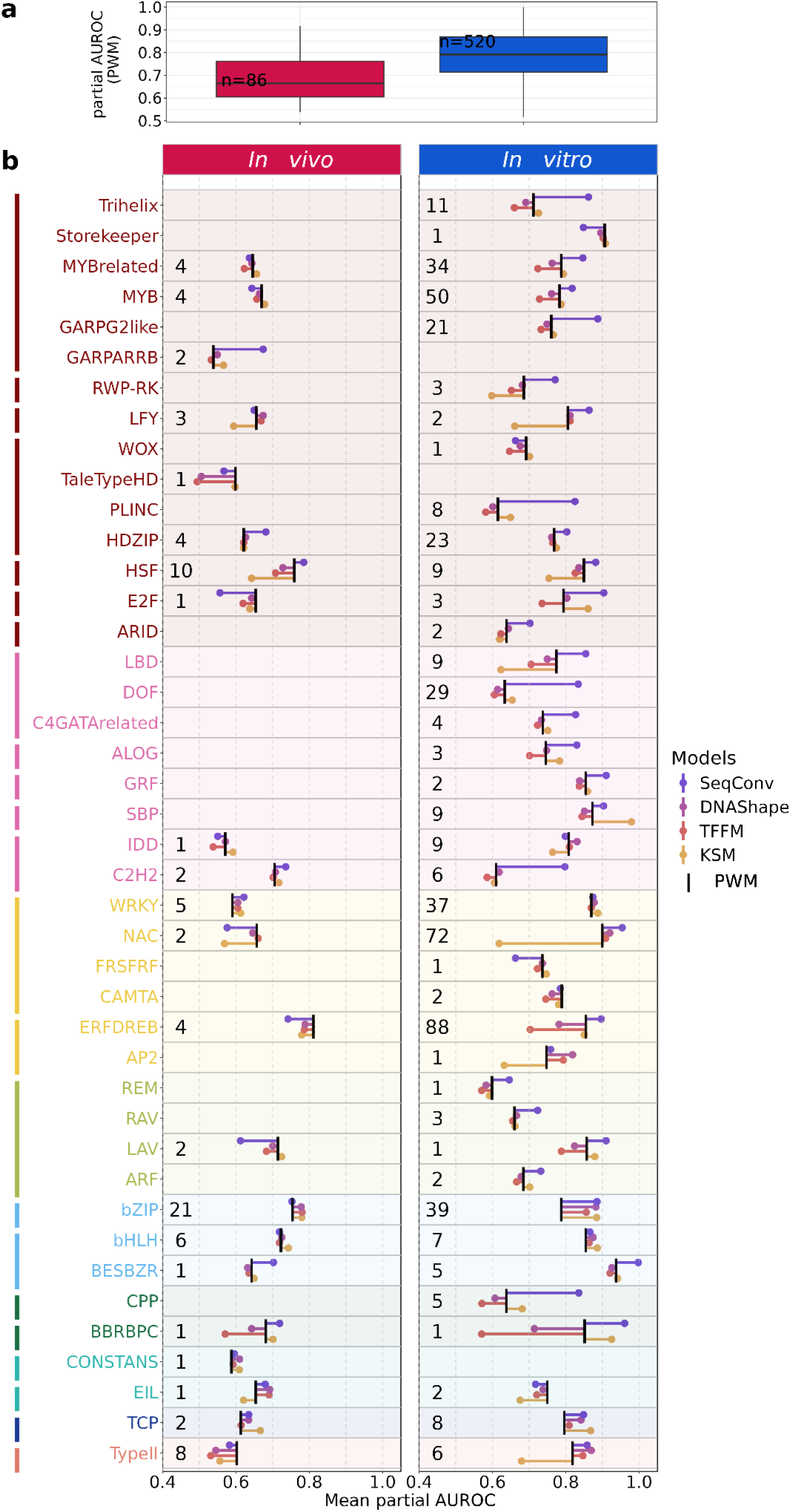
Performance of transcription factor binding models across families evaluated at low false-positive rates. Same analysis as in Fig. 2, except that model performance was evaluated using the partial area under the receiver operating characteristic curve (pAUROC) restricted to the low false-positive-rate region (FPR ≤ 1%). pAUROC values were standardized to range from 0.5 (random prediction) to 1.0 (perfect prediction). **a**, Distribution of PWM performance across all TF binding datasets for *in vivo* (ChIP-seq) and *in vitro* (DAP/ampDAP-seq) experiments. **b**, Mean pAUROC per TF family for *in vivo* (left) and *in vitro* (right) datasets. Families are colored by structural superclass and ordered by TF class along the y-axis; numbers indicate the number of datasets per family. PWM is shown as the reference model (black tick), and alternative models (TFFM, KSM, DNAshape and SeqConv) are represented as lollipops indicating the deviation in mean pAUROC relative to PWM. Positive values indicate improved performance and negative values indicate reduced performance. Values were logit-transformed prior to family-level averaging and back-transformed for visualization.

**Supplementary Fig. 5:**
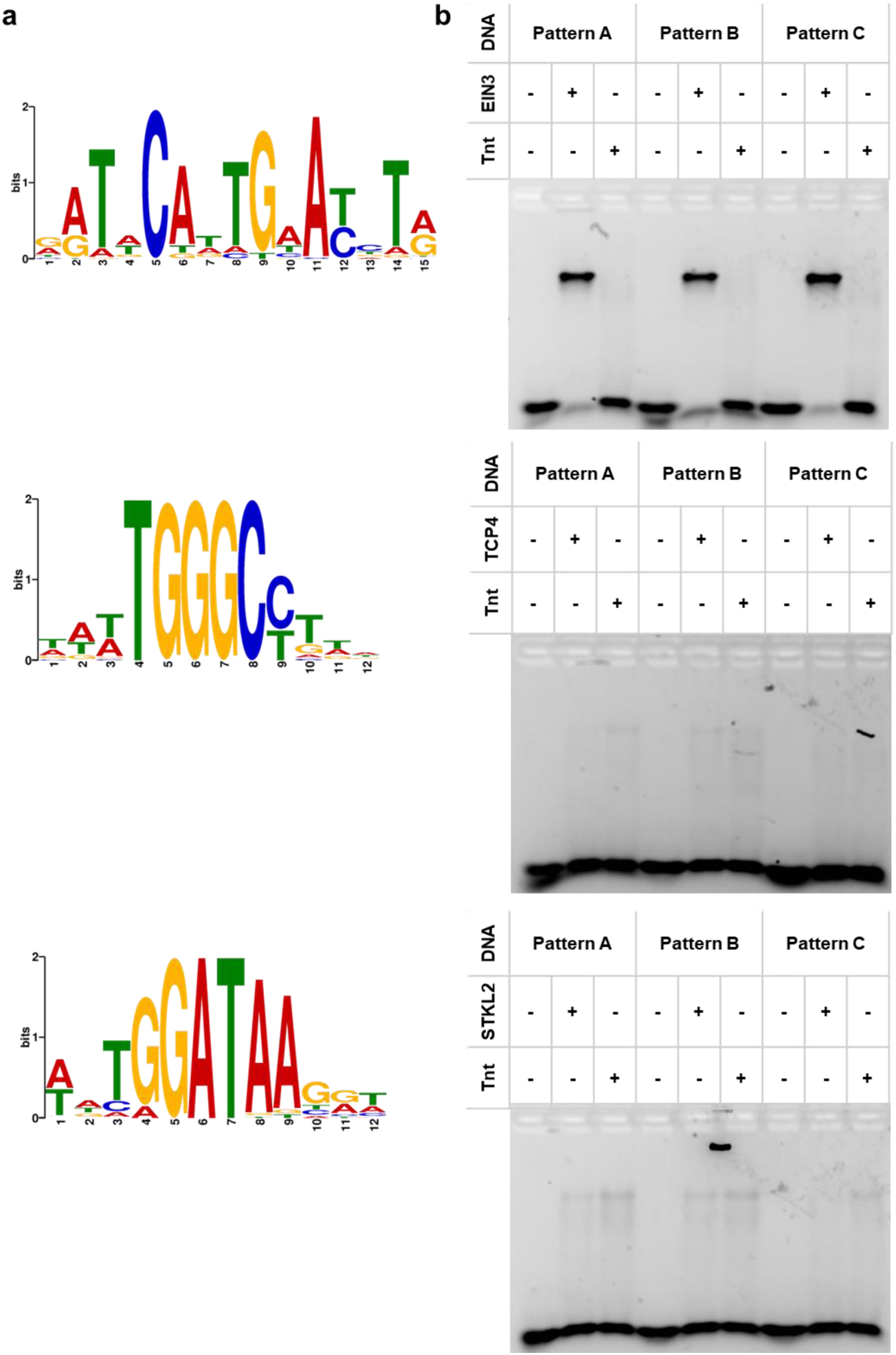
*in vitro* validation of candidate transcription factor binding motifs. **a**, Sequence logos of the candidate binding motifs selected for electrophoretic mobility shift assay (EMSA). Candidate motifs were identified during manual curation of the atlas for EIN3, TCP4 and AtSTKL2. **b**, EMSA performed with recombinant EIN3, TCP4 and AtSTKL2 proteins expressed *in vitro*. Three DNA probes corresponding to the candidate motifs of each transcription factor (Supplementary Table 7) were tested. A mobility shift was detected only for the canonical EIN3 motif (Pattern B), whereas no detectable binding was observed for the TCP4 or AtSTKL2 probes under the conditions tested.

**Supplementary Table 1: 1,657 *Arabidopsis thaliana* TFs classified in 56 Plant-TFClass families.**

**Supplementary Table 2: Binomial tests for positional bias of significantly enriched TFBS–TSS distances.**

**Supplementary Table 3. Binomial tests for orientation bias of TFBS at significantly enriched TFBS–TSS distances.**

**Supplementary Table 4. Conservation of homotypic spacing enrichments between *in vitro* and *in vivo* datasets for transcription factors represented in both conditions.**

**Supplementary Table 5. Complete metadata and processing parameters for all ChIP-seq, DAP-seq, and ampDAP-seq datasets analyzed in this study.**

**Supplementary Table 6. Primers used for cloning transcription factors for EMSA validation.**

**Supplementary Table 7. DNA probes used for EMSA validation of candidate transcription factor binding motifs.**

**Supplementary Table 8. Summary of TF-specific TFBS positioning preferences relative to transcription start sites.**

**Supplementary Table 9. Summary of homotypic TFBS spacing configurations detected across Arabidopsis transcription factors.**

*Supplementary tables are in the associated “supplementaryTables.xlsx” file*.

